# The digestion time for salmon louse (*Lepeoptheirus salmonis*) in lumpfish (*Cyclopterus lumpus)* in relation to freshness, developmental stage, and temperature

**DOI:** 10.1101/2024.09.14.613060

**Authors:** Kirstin Eliasen, Sandra L. Østerø, Tróndur T. Johannesen, Esbern J. Patursson, Ása Jacobsen, Agnes M. Mortensen, Marner Nolsøe, Ása Johannesen

## Abstract

Sea lice infestations cause significant economic losses in the Atlantic salmon aquaculture industry. To biologically control sea lice at farming sites, cleaner fish such as lumpfish are employed. However, the efficacy of lumpfish is under constant debate, primarily due to limited knowledge of digestion times, which makes it difficult to interpret the number of salmon lice found in the stomach contents of dissected lumpfish.

The aim of this study was to provide quantitative estimates of the degradation of salmon lice over a period of 12 days. After an acclimation period of approximately one week, batches of eight lumpfish (average weight 94.3 g, SD ± 33.2) were fed salmon lice and arranged in tanks.

Each batch received six large mobile lice and two adult female lice. Samplings were conducted at 24-hour intervals during the first four days and at 48-hour intervals over the remaining eight days. The experiment was conducted twice, each at a different temperature regime (6°C and 9°C), using live lice in both trials. To investigate if the freshness of the louse influenced degradation and digestion, the setup was replicated in the 9°C experiment with lice that had been stored frozen at -80°C, with an additional 12-hour sampling point for comprehensive observation.

The analysis of salmon lice revealed expected digestion times of 6.8 days and 12 days for large mobile and adult female salmon lice, respectively. Temperature and lice freshness did not seem to influence digestion times, but the developmental stage of the lice did. The findings of this study can be used to estimate the cleaning efficacy of lumpfish based on the stomach contents.

## Introduction

Sea lice have posed a serious problem for the Atlantic salmon farming industry since the 1970s [1], with their economic impact surpassing that of any other farmed salmon parasite [2]. The increasing resistance of sea lice to medical treatments necessitated the exploration of alternative, non-pharmaceutical methods [3–6], and as a result, the use of cleaner fish emerged as a method for controlling sea lice [7].

Several fish species have been identified as effective cleaners, particularly among the wrasses [8–9]. However, the wrasse species currently used for biological delousing are temperature-sensitive, making them unsuitable for use at low temperatures [10]. In contrast, lumpfish (*Cyclopterus lumpus*) have proven to be effective cleaners at lower temperatures [11–12]. Due to the relatively low temperatures and the absence of native wrasse species, lumpfish is the only cleaner fish used in the Faroe Islands [13].

Since the onset of using lumpfish as cleaner fish, their cleaning effort has shown to vary greatly, both spatially, seasonally and with size [11, 13–17]. According to Staven et al. [17], there are two primary approaches to assessing the cleaning efficacy of lumpfish. The first method involves indirectly estimating efficacy by comparing sea lice infestation levels in cages with and without lumpfish. This approach has been demonstrated in controlled experiments [12, 18] and modelling studies [19]. However, both methods are significantly influenced by the complexity of sea lice population dynamics at both the site and on cage level [15]. The second, more direct method involves reviewing data and personal experiences from fish farmers, as conducted by Imsland and Reynolds [20].

An alternative approach to measuring cleaning efficacy is to investigate and count the presence of lice in the stomach contents of lumpfish, as has been done in numerous studies [11, 13, 15–16]. However, this method must be combined with assumptions about digestion time, underscoring the importance of understanding the digestion time for salmon lice in lumpfish. Additionally, the use of cleaner fish for biological control has raised ethical considerations, particularly regarding the welfare and high mortality rates frequently observed when cleaner fish are stocked in sea cages [21–23]. Given these concerns about the efficacy and welfare of cleaner fish in commercial sea cages, forceful evidence is required to justify and guide their use in the industry.

Here, we investigate the digestion times of salmon lice in lumpfish stomachs primarily to assess the effects of temperature and louse life stage. Salmon lice were inserted into the stomachs of live lumpfish and left to incubate for predetermined time intervals. Following incubation, the lumpfish were dissected to determine the digestion status of the lice.

## Material and methods

### Ethical statement

The use of lumpfish for experimental purposes was accepted by the Faroese Food and Veterinary Authority (HFS 23/0331-5 and 23/00331-11). Additionally, Firum’s Animal Experimentation Ethics Committee reviewed and approved the study (approval number 2023_01). The committee’s decision was based on the anticipated welfare benefits for lumpfish in aquaculture and the minimal suffering projected to result from the study.

Lumpfish were sourced from commercial salmon farms and transferred to the experimental facilities in tanks with aerated seawater by car. All fish were housed in communal tanks, fed ad libitum throughout the study, and provided with shelters. Sedation was performed in dark buckets with aerated seawater using a low dose of MS-222 (100 mg/L, according to Skår et al. [24]) until the fish showed mild signs of sedation (poor balance) to minimize stress during the experimental procedure. Euthanasia was carried out with an overdose of MS-222 (1 g/L, according to Skår et al. [24]) for a minimum of 15 minutes, after which the fish were dissected immediately to ensure they did not regain consciousness.

Any fish showing signs of poor welfare, such as injuries or abnormal behaviour, would be euthanized before the end of the trial. However, all the fish in this study were healthy and did not warrant any intervention.

### Research animals

The lumpfish used in this study were collected from salmon cages at a Hiddenfjord farming site, one to two months post deployment. The fish originally came from Benchmark Genetics Iceland hf in Iceland, where the lumpfish broodstock used to make the hatchery reared fry were wild caught. The lumpfish were transported to the Faroe Islands in marine shipping containers. The lumpfish were fed to satiation on commercial feed pellets (Skretting, Clean Lumpfish 3, 3 mm), which was the same feed as the lumpfish were fed at the farming site, once or twice daily throughout the duration of the experiment.

At arrival, the lumpfish (*n* = 208 split in two rounds of trials with 144 and 64 lumpfish each) were distributed among experimental housing tanks. The tanks were white-bottomed, black-sided fiberglass tanks with a capacity of 125 litres, measuring approximately 50 cm × 50 cm × 50 cm. Maximum biomass in each tank was 10 kg/m^3^. All tanks had a flow-through system with aerated seawater, maintaining a flow rate of two litres per minute, equating to a full tank exchange every hour. Each tank was equipped with shelters made of black PE drainage pipes cut in half lengthwise and hung vertically in pairs (40 cm long, with a total area of 0.12 m²). The overhead lights were set to a 12:12 light:dark schedule.

To ensure that any salmon lice consumed at the farming site did not affect the study results, the lumpfish were not fed a salmon louse until seven to nine days after their arrival at the research facility. By the end of the experiment, the lumpfish had a mean weight of 94.3 g, with a standard deviation (SD) of 33.3 g, reflecting the size of fish in the early phase of deployment.

Salmon lice used in the study were collected from salmon farms during the mandated sea lice counts, with the collection occurring within days before the start of the trial. The lice were sampled directly from the salmon, placed in buckets filled with seawater, and transported by car to the research facility. To ensure the lice remained alive until the experiment began, aeration was provided to the buckets, and the water was maintained at in situ temperatures. However, for comparative purposes, a fraction of the sampled salmon lice was frozen at -80°C immediately upon arrival at the research facility.

### Experimental setup and sampling

The study consists of three treatments (freshness, developmental stage and temperature) divided into two trials conducted at separate times. The first trial compared digestion time of frozen and live salmon lice where half the lumpfish were fed live lice and the other half were fed frozen lice. Additionally, to compare digestion time of developmental stage of lice, one quarter of the lumpfish (two out of eight fish per sampling point) were fed an adult female salmon louse, whereas, the rest were fed a large mobile salmon louse. Subsequently, the same trial was carried out at a lower temperature, but using only live salmon lice. The decision to not use frozen lice in the second trial was made in order to minimise the use of animals as the initial trial had clear enough results for frozen lice and establishing the effect of temperature on digestion of live lice had a higher priority.

To facilitate handling, lumpfish were anesthetized using a solution containing 100 mg/L MS-222 (Tjaldurs Apotek, Tórshavn, Faroe Islands) until they exhibited signs of sedation, such as cessation of swimming, loss of equilibrium, and lack of responsiveness, in accordance with the protocol outlined by Skår et al. [24]. Subsequently, and in batches of eight, the sedated lumpfish were administered a single salmon louse via oral insertion, guiding it past the oesophagus using rounded-tipped forceps. Within each batch, two lumpfish were administered an adult female salmon louse, while the remaining six were fed a large mobile salmon louse, i.e. a preadult II or adult male salmon lice (52 adult females and 156 large mobile).

After receiving the salmon lice, each batch of lumpfish (*n* = 8) was assigned to a designated tank (18 tanks in total), representing sampling points at intervals of 0.5, 1, 2, 3, 4, 6, 8, 10, or 12 days after the feeding event (*n* = 72 for frozen lice and live lice, respectively). All lumpfish were also offered their usual lumpfish feeds throughout the incubation period to ensure that digestion activity remained normal and like that at a farming site. The second trial did not include the 0.5-day sampling point, as all lice were recovered after 24 hours at the warmer temperature in the previous trial (*n* = 64, eight tanks). The selection of sampling points was based on a previous study, which found that 66% of adult male salmon lice could be visually identified three days after being consumed by lumpfish [25]. All fish recovered seamlessly from the sedation, displaying no signs of distress or adverse effects from the procedure, and there were no recorded mortalities throughout the experiment.

Temperature readings were recorded using RBR Solo^3^ D temperature loggers, capturing data at 10-minute intervals. The experiment was conducted twice, each at a different temperature regime: first, from December 9, 2023, to December 21, 2023, with an average temperature of 9.01°C (minimum: 7.64°C, maximum: 9.52°C) and with an average lumpfish weight of 89.7 g, with a standard deviation (SD) of 20.9 g; and second, from March 1, 2024, to March 13, 2024, with an average temperature of 6.25°C (minimum: 5.14°C, maximum: 6.68°C) and with an average lumpfish weight of 104.5 g, with a standard deviation (SD) of 49.7 g.

At each designated sampling point, following the protocol established by Skår et al. [24], lumpfish underwent euthanasia through a 15-minute exposure to a 1 g/L MS-222 solution. Subsequently, their stomachs were carefully dissected for content assessment. The measurements conducted included time elapsed since feeding, weight, length, and determination of sex. Stomach contents were categorized based on the presence of (1) adult female salmon lice, (2) mobile stage of salmon lice, and (3) lumpfish feed. Furthermore, each salmon louse discovered was evaluated for signs of degradation, with categorizations of (0) no apparent degradation, (1) slight signs of degradation, and (2) significant signs of degradation. A rating of 1 indicated partial digestion of the soft tissue, while a rating of 2 signified complete digestion of the soft tissue (Fig 1).

**Fig 1.**
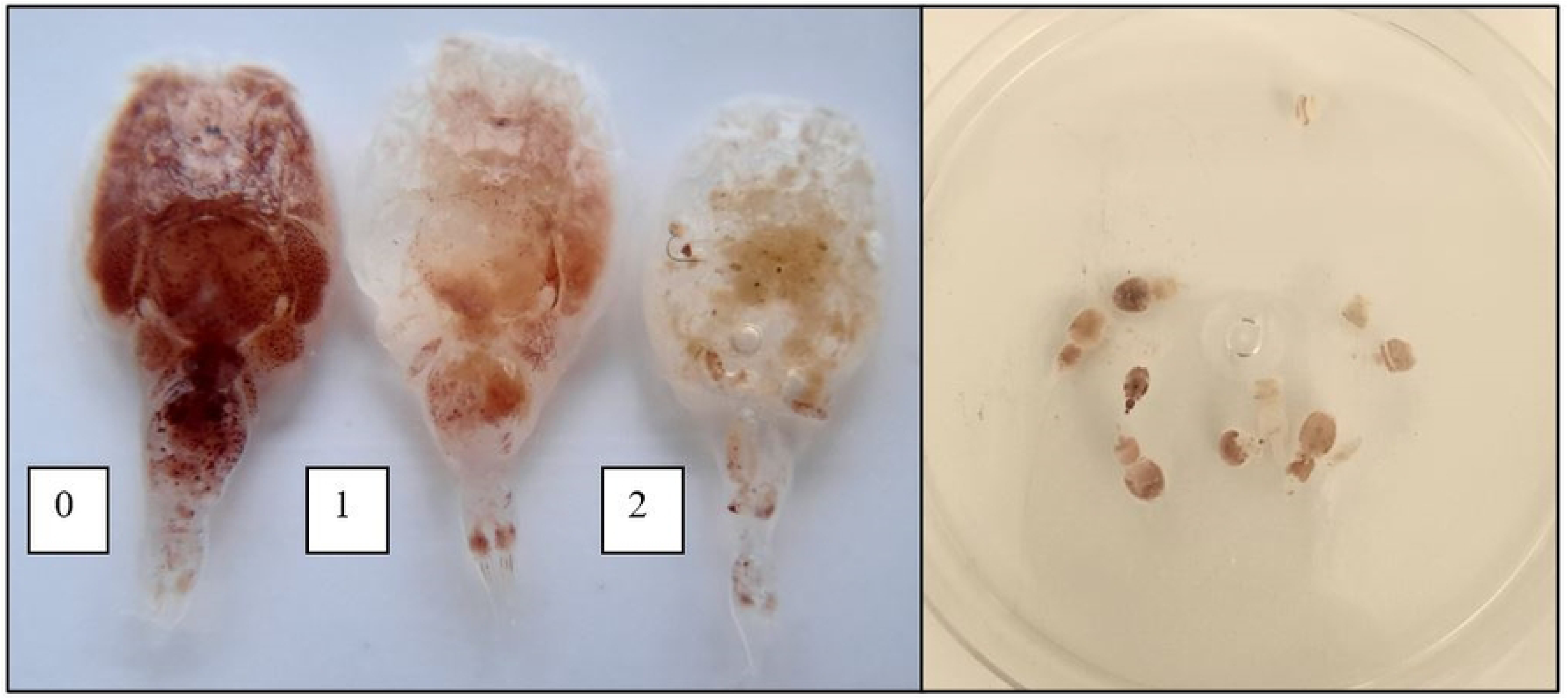
Degradation categories for salmon lice were defined as follows: (0) no apparent degradation, (1) slight signs of degradation, and (2) significant signs of degradation (left). The 11 adult female salmon lice and one mobile salmon louse found in a lumpfish stomach at four days incubation time (right).

Despite being housed at the research facility for at least a week prior to the commencement of the experiment, one lumpfish, which had received a frozen louse, was discovered with 11 adult female salmon lice and one mobile salmon louse in its stomach at four days incubation time, making it impossible to determine the developmental stage of the louse and its degradation category. This occurrence suggests that the lumpfish retained ingested salmon from the farming site in its stomach for at least 11 days after ingestion, but the fish had to be excluded from the data analysis (*n* = 207).

### Statistical analysis

Survival analysis was used for estimating the probability of recovering salmon lice over time. The response variable was binary (i.e., identified, or unidentified) at a given time, and the data were a mix of left- and right-censored data, indicating that the true identification times were unknown and could have occurred either before or after a sampling point. The analysis was done in two stages: A non-parametric survival analysis was done to provide an overview of survival probabilities using a Kaplan-Meier estimator. Afterwards, a parametric survival regression using the Weibull distribution was performed to model the survival probabilities over time.

First, a combined survival regression model was fitted to assess the overall significance of developmental stage, temperature, and freshness as predictors. Secondly, the nonsignificant predictors were excluded from the model, and separate regressions for the two developmental stages was fitted. This approach allowed for different scale parameters for each developmental stage, which captured the characteristics of each stage better.

The analysis was performed using the survfit and survreg functions in the survival package in R (www.r-project.org) [26]. For estimation of mean digestion times, the survival curves were integrated.

To determine factors influencing the degradation levels of salmon lice, a cumulative link model (CLM) analysis was used to investigate the interaction between time and freshness (frozen vs live lice), temperature and developmental stage on degradation level. The model included degradation level as ordinal response, and interaction terms between time and each of the factors.

The CLM was fitted using the clm function from the ordinal package in R (www.r-project.org) [27].

In all statistical tests, results were considered significant at p < 0.05.

## Results

### Salmon louse recovery and degradation over time

The proportion of salmon lice recovered and the grade of degradation of the lice over time since feeding is illustrated in Fig 2. All lice were recovered within 24 hours. After two days, only the frozen lice were not fully recovered. Notably, lice were still present in the lumpfish stomachs after twelve days.

**Fig 2.**
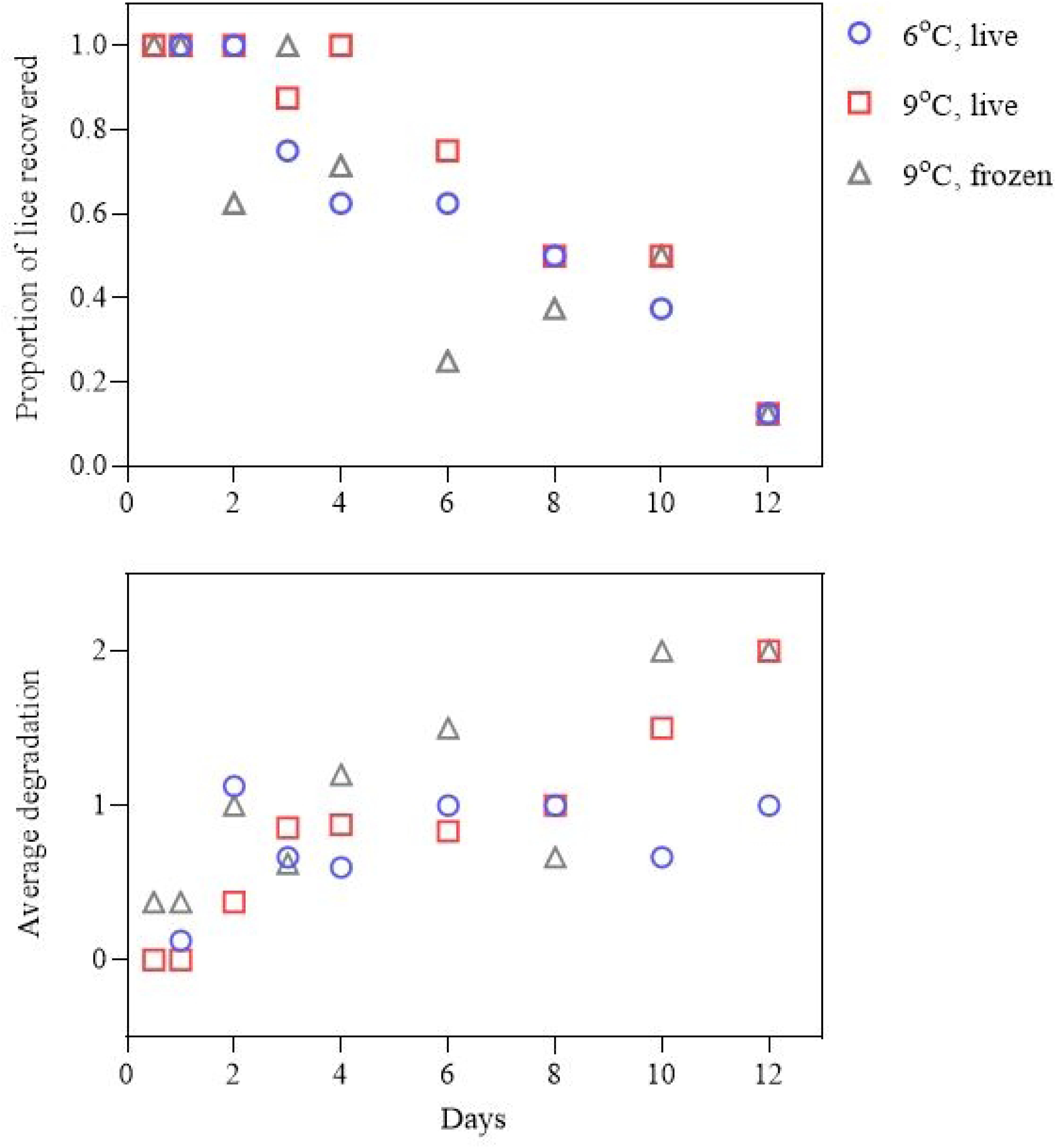
Proportion of lice recovered (upper) and average salmon lice degradation (lower) for the various times since feeding. The observations are coloured and shaped by temperature and freshness (frozen vs. live).

### Factors influencing salmon louse recovery

The survival regression analysis showed no significant difference in the recovery of sea lice between the two temperatures (deviance_1,204_ = 0.121, p = 0.728) or freshness (deviance_1,204_ = 1.692, p = 0.193). These p-values indicate that neither temperature nor freshness (frozen or alive) significantly influenced the recovery rates of sea lice. The analysis showed a significant difference (deviance_1,204_ = 19.268, p < 0.001) in the proportion of lice recovered between the two developmental stages of salmon lice. The probability of recovering a mobile salmon louse thus decreased significantly faster over time compared to adult female salmon lice (Fig 3).

**Fig 3.**
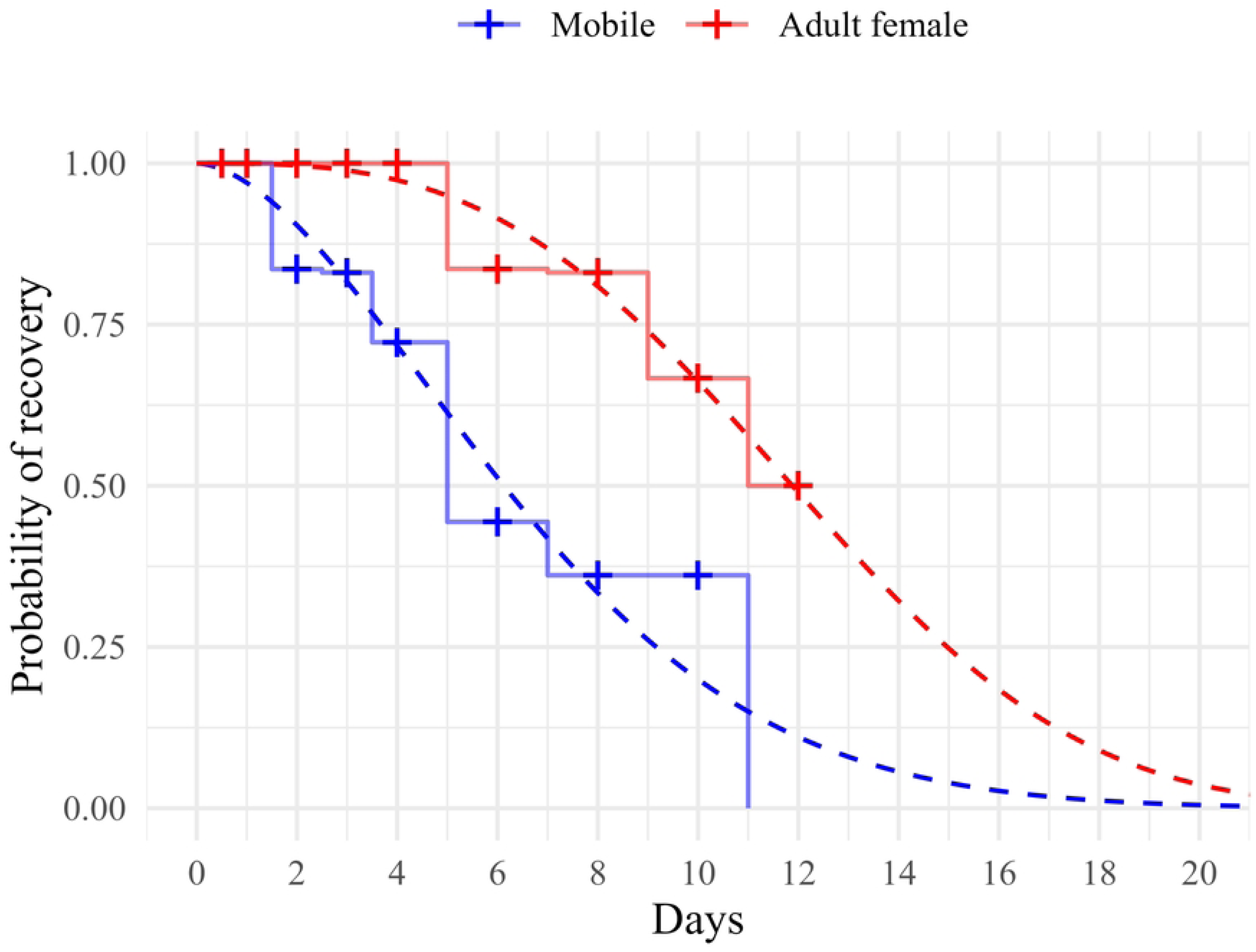
The probability of recovering adult female and mobile salmon lice from lumpfish stomachs across different sampling times. Full lines are from the non-parametric survival analysis, and broken lines are from the fitted parametric survival regression. ‘+’ indicates sampling points.

All lice were recovered within the first day. However, from the second day onward, the proportion of mobile salmon lice steadily decreased, and by day 12, no mobile lice were recovered. In contrast, all adult females were recovered until day four, and by day 12, half of the adult females were recovered.

Since the lumpfish were fed salmon lice in batches without individually tagging the lumpfish, the developmental stage of the lice consumed by each lumpfish was unknown. This lack of information about the lice’s developmental stages, which significantly affected digestion times, impeded the investigation into how other factors, such as lumpfish sex and size, influenced the digestion rate.

### Expected digestion times

Integrating the survival curves resulted in the expected digestion times of 6.8 days for large mobile salmon lice and 12 days for adult female salmon lice.

### Factors influencing salmon louse degradation

The interaction between time and developmental stage was significant (z = -2.976, n = 207, p = 0.003), indicating that mobile lice degraded faster than adult female lice. No other significant interactions were found, indicating that temperature (z = -1.084, n = 207, p = 0.278) and freshness (z = 0.710, n = 207, p = 0.477) did not affect degradation over time. However, there was a main effect of temperature (z = 2.119, n = 207, p = 0.034), indicating more rapid degradation at the start of digestion, and freshness (z = -3.133, n = 207, p = 0.002), indicating that frozen lice start out more degraded than live lice.

## Discussion

Understanding the digestion time of salmon lice in lumpfish is crucial for assessing their cleaning efficacy. This knowledge enables the estimation of number of salmon lice consumed per unit of time based on the number of salmon lice found in stomach contents [15]. Given that the welfare of lumpfish is frequently compromised in sea cages [21–23], obtaining accurate information on the digestion time of salmon lice in lumpfish is urgently needed. This study provides essential insights, offering robust guidance for further use of lumpfish in the aquaculture industry.

To the authors’ knowledge, this is the first study to investigate the probability of recovering salmon lice fed to the lumpfish alive as a function of time since feeding. While Eysturskarð et al. [25] also used live salmon lice, their sampling period was too short - only three days - to accurately estimate digestion time. In contrast, Staven et al. [17] conducted a comparable, longer-duration experiment, but with the use of frozen lice.

Our findings on the digestion time for salmon lice in lumpfish (6.8 and 12 days) differ significantly from those of Staven et al. [17], who estimated a digestion time of 29 hours at 9°C. As the results from Staven et al. [17] were published while our experiments were still ongoing and it was already apparent that our results deviated substantially, we decided to conduct an additional experiment using frozen lice. However, no significant difference was observed in digestion time between frozen and live lice, which aligns with the findings of Jackson et al., who found no significant difference in the digestibility of four types of prey whether they had been frozen or not [28].

In the Faroe Islands, salmon lice are found in approximately 3.6% of the lumpfish stomachs examined (unpublished data, n = 26,036), while in Norway, the frequency is slightly lower at 3.1% [16]. The average number of salmon lice per stomach is also higher in the Faroe Islands, with 0.41 lice per stomach (unpublished data, n = 26,036), compared to 0.19 lice per stomach in Norway [16]. Assuming a relatively equal cleaning efficacy in the two locations, the difference in lice numbers is not sufficient to explain a several fold difference in digestion time. Moreover, Engebretsen et al. attributed the difference in lumpfish lice consumption between the two locations to a higher threshold level of salmon lice in the Faroese regulations compared to the Norwegian regulations [16].

Another difference between this experiment and that of Staven et al. [17] is the feeding regime. In the current experiment, the lumpfish were fed lumpfish feed once or twice a day throughout the duration of the experiment to simulate the conditions typically experienced by lumpfish used as cleaner fish. In contrast, in the study by Staven et al., the lumpfish did not receive any food after being fed the salmon lice [17]. Andersen [29] showed how the stomach evacuation time of whiting increased when the fish were fed an increasing number of herring, and it is thus plausible that the digestion time for the salmon lice could be dependent on the overall diet of the lumpfish. However, in the Eysturskarð et al. experiment, lumpfish were not given any feed after consuming an adult male salmon louse, yet 66% of the lice were still visually detectable on day three [25].

In 1990 Hopkins and Larson studied the gastric evacuation of three food types in the black and yellow rockfish (*Sebastes chrysomelas*). Here they found digestion times to be dependent on the food type. When the rockfish were fed fish, an intrinsic digestion pattern, characterized by a period of rapid evacuation followed by a less rapid decline, typically occurring when small friable foods of low energy are ingested, was observed. When the rockfish had ingested shrimps or crabs, the observed pattern was a linear decrease with time, but with an initial lag phase [30]. Hopkins and Larson suggested that the lag phase observed was due to the hydration of chitin, which is an enzymatic reaction requiring acidic conditions for the digestion of a chitinous prey to begin. The secretion of chitinase and acid, and the time required until the exoskeleton of the food can be digested or softened sufficiently to be disrupted by gastric musculature, was thus suggested to account for the lag phases observed [30].

The digestion patterns observed in present study are comparable to those observed when the rockfish were fed shrimps or crabs, i.e. with an initial lag phase (Fig 2). However, the duration of the lag phase was significantly longer regarding the digestion of adult female salmon lice, compared to the lag phase of the mobile salmon lice digestion (Fig 3). On the contrary, the digestion pattern observed in Staven et al. was more comparable to the intrinsic digestion pattern observed in rockfish when fed fish, i.e. having a period of rapid evacuation followed by a less rapid decline [17]. However, in Staven et al. some lumpfish were also observed to have salmon lice in their stomach three to seven days after ingestion. Unfortunately, two salmon lice were found in the filters of their experimental tanks, proving regurgitation of lice, and the authors thus speculated that these outliers could be lumpfish that had ingested regurgitated lice on a later day [17]. However, the lumpfish which still had salmon lice in its stomach on day three, had four lice in its stomach, and the lumpfish which had salmon lice in its stomach on day seven, had two salmon lice, indicating a substantial number of regurgitated lice. If ingestion of regurgitated lice on a later date was the case, salmon lice not recovered at an earlier date could be flawed data as well.

Another key difference between the study by Staven et al. [17] and the present study is the origin of the lumpfish. In the latter experiment, the lumpfish were derived from wild-caught broodstock. In contrast, the lumpfish used in Staven et al. originated from a breeding program where the roe came from wild-caught females, and the milt was collected from captive male broodfish belonging to the broodstock nucleus of Namdal Rensefisk AS and AquaGen AS, specifically the Namdal Rensefisk AS GEN2 selected strain. This breeding process suggests that certain characteristics might have been selectively bred into the lumpfish used in Staven et al. [17].

While lumpfish are generally considered omnivorous feeders [13, 31], Imsland et al. found that different families of lumpfish have preferences for different types of food. The family that most frequently ingested sea lice was also the one that actively sought out natural food sources, such as crustaceans, over available feed pellets, resulting in more rest periods compared to other families. The authors speculated that this behaviour could be genetically based [32]. The findings of sea lice and crustaceans in the same lumpfish in Imsland et al. [32] align with other findings showing a positive association between the prevalence of organisms associated with biofouling, e.g., the crustaceans *Caprella* spp. and *Tubularia* spp., and sea lice in the stomachs of lumpfish sampled from salmon pens [13, 16].

If food preferences are genetically based, it is possible that digestive traits, such as the ability to digest chitinous prey faster due to these food preferences, are inheritable as well. If the cleaning trait has been actively bred into the lumpfish used in Staven et al. [17], these fish might be able to digest salmon lice faster than the lumpfish used in the present study. If survival in salmon pens depends on the ability to digest chitinous prey quickly, selection for this trait could potentially influence digestion time as well. However, if this is the case, estimating the cleaning efficacy of lumpfish used as cleaner fish becomes even more challenging.

Inherited differences in the ability to digest chitinous prey could explain the frequently observed phenomenon of small, poorly lumpfish with hundreds of salmon lice in their stomachs, sampled from salmon pens, as the presence of low levels of chitinase in fish with physical prey disruption mechanisms would not be sufficient to have any significant digestive action [33]. In these cases, the enzyme may function by dissolving awkwardly shaped chitinous structures that become lodged in the gastrointestinal tract, causing a blockage. Gut blockage is a common cause of mortality, especially among young fish [33].

Numerous studies have shown that water temperatures can influence digestion rates in fish [34–38]. However, these studies are all conducted with larger temperature gaps than present study. Nevertheless, no significant difference was found in the digestion rate for salmon lice in lumpfish at 6°C and 9°C (Fig 2).

The significant difference in digestion times for adult female and large mobile salmon lice suggests that lumpfish might forage on a larger proportion of salmon lice at lower developmental stages than previously estimated based on stomach content analysis. In the Faroe Islands, the adult female to mobile salmon louse ratio in lumpfish stomachs sampled from salmon pens is approximately 1:1.2 (unpublished data). However, when accounting for the difference in digestion time, this ratio adjusts to approximately 1:2.1, which aligns more closely with the adult female to mobile salmon louse ratio on the salmon, typically varying between 1:2 and 1:4.2 [39]. This finding indicates that lumpfish might not be as selective in their foraging on adult female salmon lice as originally suggested [11–12].

Overall, the reasons for the significant differences observed between the results of the current study and those of Staven et al. [17] should be investigated in future research. Potential areas of investigation should include examining within-species differences in digestive traits, such as pH and chitinase levels. Although neither study stimulated physical activity in the lumpfish, movement has been shown to circulate digestive enzymes [28], and the influence of physical activity on digestion time for salmon lice in lumpfish should be included in future studies.

Additionally, while no significant difference was found in digestion times at 6°C and 9°C in this study, the temperature difference might have been too small to detect potential temperature-induced variations. Future studies should aim to investigate digestion times at a wider range of temperatures to better understand the influence of temperature on the digestion process.

## Conclusions

To deduce the cleaning efficacy from the number of lice found in lumpfish stomach samples, it is essential to know the digestion time of salmon lice in lumpfish. In this study, the expected digestion time for salmon lice in lumpfish was determined to be 6.8 days for large mobile lice and 12 days for adult female lice, at temperatures ranging from 6°C to 9°C. These results differ substantially from previously published findings [17]. While Staven et al. [17] suggested that the estimated number of salmon lice consumed per lumpfish per day could be calculated by dividing the number of lice found in a lumpfish stomach by 1.39, our results indicate a significantly lower cleaning efficacy. According to our findings, the number of retrieved salmon lice should be divided by 6.8 for large mobile lice and 12 for adult female lice to estimate the number of salmon lice consumed per lumpfish per day.

## Acknowledgments

The authors would like to thank Heri Christoffersen and his staff at the farming site, as well as Unn V. Johannesen, Jógvan F. Hansen and Sverre Muller at Firum for assistance and technical support.

